# An early Shh-H_2_O_2_ feedback loop controls the progression of the regenerative program during adult zebrafish fin regeneration

**DOI:** 10.1101/2021.08.19.456615

**Authors:** Marion Thauvin, Rodolphe Matias de Sousa, Marine Alves, Michel Volovitch, Sophie Vriz, Christine Rampon

## Abstract

Reactive oxygen species (ROS), originally classified as toxic molecules, have attracted increasing interest given their actions in cell signaling. Among these molecules, Hydrogen peroxide (H_2_O_2_) is the major ROS produced by cells and acts as a second messenger to modify redox-sensitive proteins or lipids. After amputation, tight spatiotemporal regulation of ROS is required first for wound healing and later to initiate the regenerative program. However, the mechanisms carrying out this sustained ROS production and their integration with signaling pathways are still poorly understood. We focused on the early dialog between H_2_O_2_ and Sonic Hedgehog (Shh) during fin regeneration. We demonstrate that H_2_O_2_ controls Shh expression and that Shh in turn regulates the H_2_O_2_ level via a canonical pathway. Moreover, this tightly controlled feedback loop changes during the successive phases of the regenerative program. Dysregulation of the Hedgehog pathway has been implicated in several developmental syndromes, diabetes and cancer. These data support the existence of a very early feedback loop between Shh and H_2_O_2_ that might be more generally involved in various physiological or pathological processes. These new findings pave the way to improve regenerative processes, particularly in vertebrates.

## Introduction

Reactive oxygen species (ROS) were initially considered to be only deleterious compounds in living organisms but were later found to contribute to physiological processes ^1– 3^. They modulate signal transduction and stress responses by acting as second messengers for redox modification of sensitive substrates ^4,5^. Living organisms possess finely regulated systems to control cellular ROS levels, including catalase, peroxiredoxin, glutathione peroxidase and superoxide dismutase ^6^. Among ROS family members, hydrogen peroxide (H_2_O_2_) is now known to be a major second messenger acting in redox signaling pathways^7^. Indeed, its lifespan is compatible with cell signaling. H_2_O_2_ is mainly produced by NADPH oxydases (NOX) complexes in the extracellular space and crosses cell membranes through aquaporin channels ^8^.

In vertebrates, regeneration of missing body parts requires three temporary overlapping modules: an immediate injury response (or early) module, a regeneration induction (or intermediate) module and a final (or late) module of blastema growth and patterning ^9^. Interestingly, a transient increase in ROS levels is required for the regenerative programs of the Xenopus tadpole tail and notochord ^10,11^, axolotl tail ^12^, adult zebrafish fin and heart ^13,14^, gecko tail ^15^, and mouse liver and inguinal fat pad ^16,17^ as well as for ear regeneration in the African spiny mouse ^18^ (for general review, see ^6^). Amputation-induced ROS production also promotes Drosophila compensatory proliferation ^19^ and is important during planarian body regeneration ^20^ and observed after hydra bisection ^21^. During zebrafish adult fin regeneration, it has been demonstrated that H_2_O_2_ induces apoptosis and MAPK activation, triggers compensatory proliferation, attracts axons and controls Sonic Hedgehog (Shh) expression ^13,22^. H_2_O_2_ is tightly regulated in time and space for at least 24 hours after amputation ^13^. This sustained production of H_2_O_2_ is specific to regeneration and is not observed in the wound healing context. H_2_O_2_ also attracts axons, and nerves control H_2_O_2_ levels in a reciprocal interaction ^22^.

Shh, first identified as a morphogen, controls cell growth, survival, homeostasis, fate and differentiation in a wide variety of cell types (for general review, see ^23^). Dysregulation of this pathway has been implicated in several developmental syndromes and cancers. Canonical activation of the Hedgehog (Hh) pathway occurs through binding to the transmembrane protein patched (Ptch). In the absence of ligand, Ptch inhibits signaling by suppressing the activity of Smoothened (Smo), a member of the G protein-coupled receptor family. Active Smo initiates an intracellular cascade that leads to the activation of Ci/Gli transcription factors that regulate the transcription of target genes ^24^. Furthermore, there are several noncanonical Hh signaling pathways ^25,26^, some of which are Smo-independent or Smo-dependent and Gli-independent. In addition to its role in development and pathologies, Hh signaling has been shown to perform important functions during tissue repair and regeneration ^27^. In zebrafish, which has emerged as a highly successful model for studying mechanisms of tissue regeneration, Hh signaling was shown to be involved in fin blastema formation and maintenance ^22,28^ and later in fin ray patterning ^29–31^.

Connections between Shh and redox balance have been identified in different contexts. Shh protects endometrial hyperplasial cells ^32^, cortical neurons ^33^ and hippocampal cells ^34^ against oxidative stress. H_2_O_2_ levels control key steps of Shh delivery *ex vivo* and change the Shh distribution and tissue patterning during zebrafish development ^35,36^. During zebrafish fin regeneration, injured nerves induce the production of large amounts of H_2_O_2_ through activation of the Shh pathway, providing an environment that promotes cell plasticity, progenitor recruitment and blastema formation ^22,27^. These discrete relationships take on their full meaning when integrated into the broader perspective of links between metabolism and gene/signaling networks ^37^. We decided to more thoroughly investigate the mechanisms and time course of crosstalk between Hh and redox signaling during zebrafish fin regeneration. We report that H_2_O_2_ levels control Shh expression and that a feedback loop in which Shh controls H_2_O_2_ production via a canonical pathway is activated very early.

## Materials and methods

### Fish care and surgery

Zebrafish colonies (AB-Tu and nacre fish) and transgenic fish *2*.*4shh:GFP:ABC#15* ^38^ (hereafter abbreviated shha:GFP) were maintained using standard methods. The animal facility obtained an agreement from the French Ministère de l’Agriculture (No. C750512), and the protocols were approved by the Ministère de l’Education Nationale, de l’Enseignement Supérieur et de la Recherche (No. 21358-2019062813438750). To maintain healthy colonies, a 14 h light/10 h dark cycle was used, and a water temperature of 28 °C was maintained. All these parameters were computer-controlled. The maximum housing density was approximately five fish per liter. Water filtration was dependent on an Aquatic Habitats stand-alone fish housing system and was performed automatically (Aquatic Habitats Inc., FL, USA). For the experiments, adult fish (5–10 months old) were anesthetized in 0.1% tricaine (ethyl m-aminobenzoate). Caudal fins were amputated at the level of the first ray bifurcation using a scalpel blade under a binocular magnifier, and the fins were allowed to regenerate for different durations.

### Pharmacological treatments

After amputation, a maximum of five adult fish were housed per tank. An NAD(P)H oxidase inhibitor (Nox-i, #BML-El395-0010) was purchased from Enzo Life Sciences (Inc. Farmingdale, NY, USA). Cyclopamine (HH-i), a Hedgehog inhibitor (#239803), and purmorphamine (Smo-A), a Smoothened agonist (#540220), were obtained from Calbiochem (San Diego, CA, USA). Hydrogen peroxide (H_2_O_2_, #H1009) was ordered from Sigma–Aldrich (Saint Louis, MI, USA). GANT61, a Gli1/2 inhibitor (Gli-I, #3191) was purchased from Tocris Bioscience (Bio-Techne, Minneapolis, MN, USA). Fish treated with dimethyl sulfoxide (DMSO) from Sigma–Aldrich (#D-8418) were used as the control group. More information is provided in Supplementary Table S1. Fish were maintained at 28 °C in the dark and were returned to light for 1 h per day for feeding and water changing.

### Enzyme activity measurement

Catalase (CAT) and superoxide dismutase (SOD) activities were measured using commercially available kits (#707002 and #706002, respectively; Cayman Chemical, MI, USA). After amputation, fins were rapidly frozen on dry ice and stored at -80 °C. For each measured enzyme and at each time point, the experiment was performed in at least triplicate with four defrosted, mechanically dissociated fins per measurement. The assay protocols were performed according to the manufacturer’s instructions.

### ROS detection

H_2_DCFDA, 2’7’-dichlorodihydrofluorescein diacetate (#287810; Calbiochem), was used to monitor the accumulation of ROS in adult zebrafish fins as previously described ^13^. We previously observed that H_2_DCFDA staining mimicked the detection of H_2_O_2_ with the genetically encoded HyPer probe ^22^.

### Quantification of regeneration

The efficiency of regeneration was quantified at 3 dpa. The surface of the blastema was measured using ImageJ and subsequently divided by the squared length of the amputation plane. The efficiency of regeneration is expressed as a percentage of the control value.

### Immunohistochemistry

Amputated fins were fixed with 4% paraformaldehyde overnight at 4 °C and used for whole-mount immunofluorescence with a rabbit anti-phospho-histone-H3 primary antibody (#SC-8656R; Santa Cruz Biotechnology, Inc., Dallas, TX, USA) to detect proliferative cells or with a chicken anti-GFP antibody (ab13970; Abcam, Cambridge, UK) to detect GFP in shh:gfp fish. More information is provided in Supplementary Table S1. Images were acquired using a Nikon 90i camera. P-H3-positive cells were counted on the fin ray and inter-ray 2 in all segments using ImageJ. GFP labeling was quantified by measuring the fluorescence intensity along a line starting at the fin border.

### Quantitative RT–PCR

Total RNA was extracted from four caudal fins per experimental point using a Monarch Total RNA miniprep kit (#T2010S; New England Biolabs, Ipswich, MA, USA) according to the manufacturer’s instructions. Five hundred nanograms of total RNA was reverse transcribed with SuperScript II reverse transcriptase (#18064022; Invitrogen, Thermo Fisher Scientific, Waltham, MA, USA) using oligo(dT) primers. Target gene expression levels were normalized to *Rpl13a* expression. Quantitative PCR was performed using a LightCycler 480 instrument (Roche Diagnostics, Rotkreuz, Switzerland) and TaqMan methodology with a fluorescent PCR probe (FAM-NFQ/MGB) (Applied Biosystems, Thermo Fisher) or a universal probe approach (Roche Diagnostics). Each sample was tested in at least triplicate. The analysis protocols were performed according to the manufacturer’s recommendations. The primers and probes used for the TaqMan assays were obtained from the Applied Biosystems bank (Assay IDs—rpl13: Dr03119261_m1; shha: Dr03432632_m1; sod1: Dr03074067_m1; sod2: Dr03100019_m1; sod3: Dr03188773_s1; cat: Dr03099088_m1).

For the universal probe approach, probes (Roche) and primers (Eurofins Scientific, Luxembourg) were used:

| Gene | Universal probe ID | Left primer | Right primer |
| --- | --- | --- | --- |
| **gli1** | 68 | ctg cag caa aga gtt cga ca | ctc cgt gga tgt gct cat ta |
| **gli2** | 39 | cct cac cca cca cag cat | cga tcg gga ttg gtg tgt |
| **gli3** | 120 | cgc tct agc tca tct aac agt gg | tgc aaa act caa agc agg ac |
| **smo** | 6 | tca gtg aca ccc act gga ac | ctg ctt cga cca tcc aaa gt |
| **ptch1** | 6 | aca aag cag ctg cca tac tag a | att tac cgg cac act ctg ct |
| **ptch2** | 6 | cac cgc cat ctt gga gtc | gga atg ctc cca tga aca ac |
| **duox** | 120 | cac cac ctt tat gtg ctc atc t | gga tca ggt aga tgt gga aac g |
| **nox5** | 43 | tgc cgc tat cgt atg aga aa | acg gtt gat cca aat aaa gtc c |

### Statistical analysis

All data were analyzed with Prism software (GraphPad, San Diego, CA, USA). Continuous variables are expressed as the means +/-SEMs. Comparisons between multiple groups were performed via one-way analysis of variance (ANOVA) followed by Tukey’s post hoc test. Comparisons between two unpaired groups were performed using Student’s t-test. The level of significance is represented as follows: * p value ≤ 0.05; ** p value ≤ 0.01; *** p value ≤ 0.001; and **** p value ≤ 0.0001. The sample sizes are given in Supplementary Table S2.

## Results

### Direct control of the H_2_O_2_ level differs between the early and intermediate modules

To better characterize the timing and origination of sustained H_2_O_2_ accumulation during adult caudal fin regeneration (**Fig. 1A**), we studied the activity and expression of two enzymes that directly control the H_2_O_2_ level. The activities of superoxide dismutase (SOD) (which catalyzes the production of H_2_O_2_ from O_2_^-^) and catalase (CAT) (which catalyzes the decomposition of H_2_O_2_ into H_2_O) were measured over time after amputation. CAT activity decreased very early after amputation, and an increase was detected at 72 hours post amputation (hpa), when the blastema had been formed (**Fig. 1B**). In contrast, high SOD activity was observed at 15 hpa, at the time of the peak H_2_O_2_ level, corresponding to the intermediate module ^13^, and then decreased after blastema formation in the late module (**Fig. 1C**). These data suggest that the balance between production and degradation regulates H_2_O_2_ homeostasis after amputation. We examined whether variations in enzyme activity depend on the regulation of expression at the transcriptional level. The zebrafish genome encodes four SOD genes (*sod1, 2, 3a* and *3b*) and one CAT gene (*cat*). The expression of *sod1, 2 and 3a+b* (collectively, *sod3*) and *cat* was measured by real-time quantitative RT–PCR during zebrafish fin regeneration (**Fig. 1D-E**). The *cat* mRNA concentration was slightly reduced after amputation and returned to the level in unamputated fish at 6 hpa, indicating that CAT expression was partially controlled via transcriptional regulation (**Fig. 1D**). In contrast, *sod1* and *sod2* levels were not affected by amputation, suggesting that variations in SOD activity do not depend on modulation of their mRNA levels (**Fig. 1E**). *sod3* mRNA was undetectable (not shown). In summary, the elevated levels of H_2_O_2_ observed during the early and intermediate modules were characterized by changes in CAT and SOD activity, respectively.

**Figure 1:**
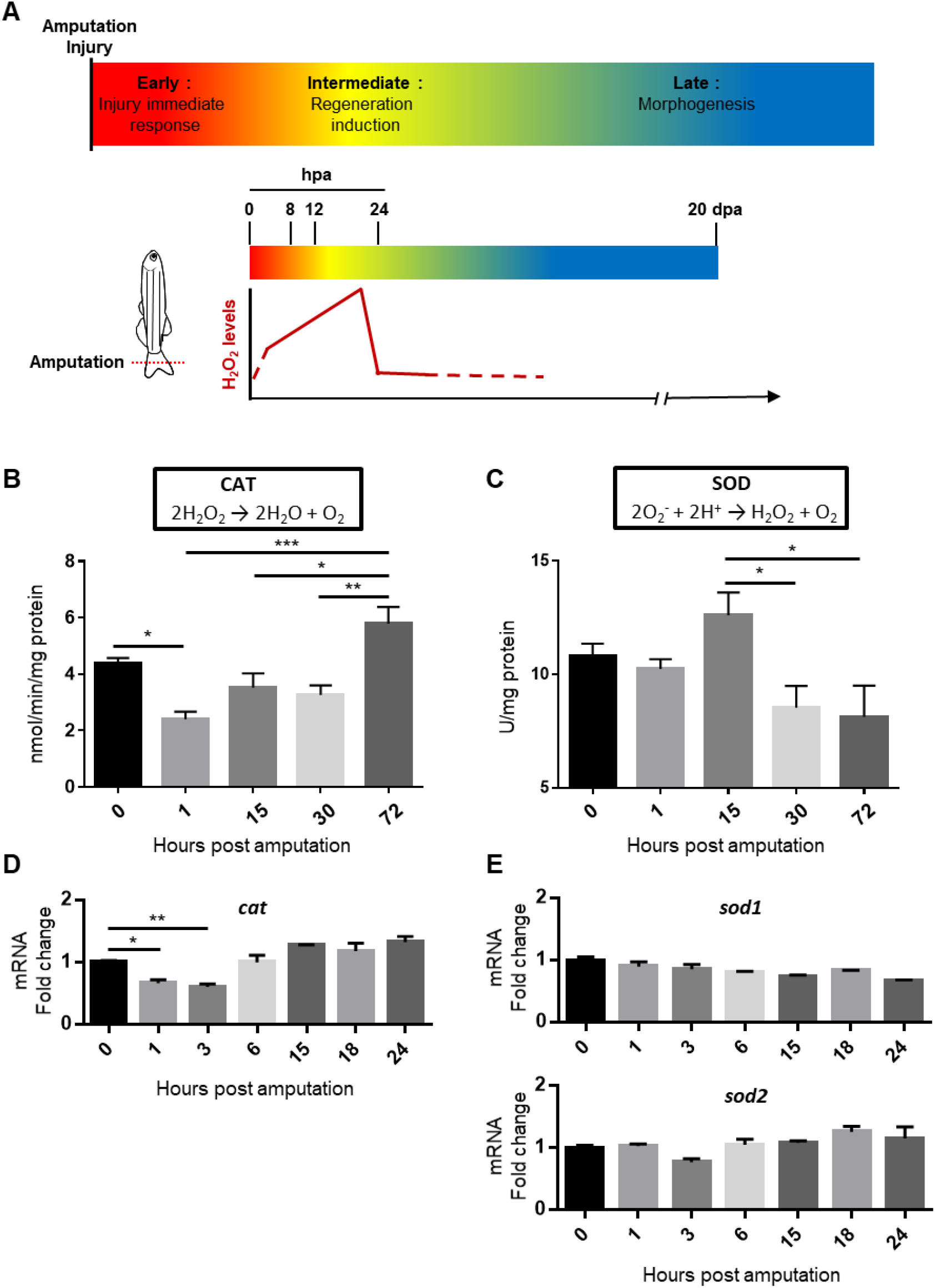
Time course analysis of CAT and SOD after amputation. (**A**) In adult tissue, regeneration is divided into three modules (upper). H_2_O_2_ levels during zebrafish fin regeneration (lower part), adapted from ^6^. (**B**) Catalase (CAT) and (**C**) Superoxide dismutase (SOD) activity levels during adult fin regeneration. (**D-E**) Gene expression was analyzed by quantitative RT– PCR during adult fin regeneration. The level in the 0 hpa sample was set to 1. The error bars indicate the SEM values (* p<0.05, **p< 0.01, ***p < 0.001). hpa: hours post amputation, dpa: days post amputation.

### Hh signaling controls SOD activity but not *sod* expression during the intermediate module

We previously showed that disruption of Smo-dependent Hh signaling with cyclopamine (Smoothened inhibitor, HH-i) treatment at 16 hpa strongly reduced H_2_O_2_ production and proliferation ^22^. To more precisely examine the role of Hh signaling in H_2_O_2_ homeostasis during the intermediate module of fin regeneration, we measured SOD and CAT activity at 3, 15 and 30 hpa in the presence of HH-i (**Fig. 2A**). HH-i strongly reduced SOD activity at 15 hpa, when the H_2_O_2_ level peaked, but had no significant effect at 3 and 30 hpa (**Fig. 2A’)**. CAT activity, as well as remaining stable over time during the intermediate module (**Fig. 1A**), was not significantly modulated by HH-i treatment (**Fig. 2A’’**). We then analyzed whether the decrease in SOD activity upon Hh signaling inhibition depends on regulation at the transcriptional level.

**Figure 2:**
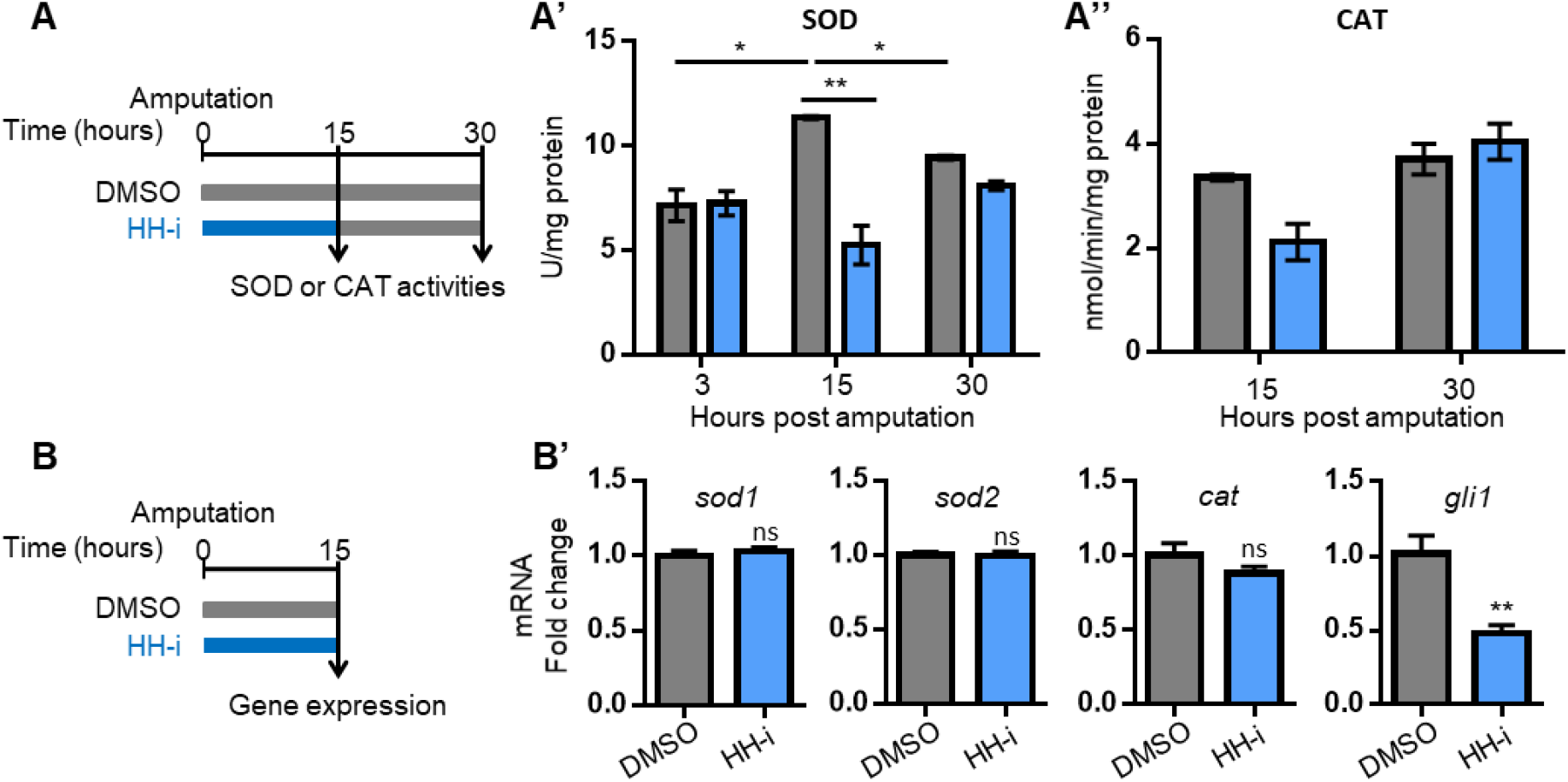
Hh signaling controls SOD activity. (**A** and **B**) Schematics of the experimental procedures. (**A’**) Superoxide dismutase (SOD) and (**A’’**) Catalase (CAT) activity was measured at 15 h (intermediate module) and 30 h (late module) post amputation after cyclopamine (HH-i) or vehicle (DMSO) treatment. HH-i was added to the water bath from 0 to 15 hpa. (**B’**) Gene expression was analyzed by quantitative RT–PCR in the regenerated fin at 15 hpa after HH-i or DMSO treatment. The level in the DMSO-treated sample was set to 1. The error bars indicate the SEM values (* p<0.05, **p< 0.01).

The mRNA levels of *sod1* and *sod2* (with *cat* and *gli1* as controls) were quantified by real-time quantitative RT–PCR at 15 hpa after HH-i or DMSO treatment (**Fig. 2B**). As expected, we observed a 2-fold decrease in the *gli1* mRNA level (**Fig. 2B’**). In contrast, the *sod1* and *sod2* (as well as *cat*) mRNA levels were not significantly affected by HH-i treatment, suggesting that HH-i reduced SOD activity without altering SOD expression.

### A feedback loop between H_2_O_2_ and Hh signaling is active during the first 24 hpa

We previously identified apparent crosstalk between Hh and H_2_O_2_ signaling during fin regeneration: inhibition of the Hh pathway strongly reduced the regeneration-specific sustained production of ROS at 16 hpa, and pan-inhibition of NOX complexes reduced the number of Shh-expressing cells at 48 hpa ^22^. This pattern was suggestive of a regulatory loop, and we decided to confirm this hypothesis by more precisely analyzing the epistatic relationship between these two signaling systems. If they were indeed engaged in a feedback loop, both should appear to act upstream of the other in the regeneration process.

We first combined NOX complexes inhibition with Hh pathway stimulation and measured the effect on stump progenitor cell proliferation at 24 hpa, a prerequisite for regeneration, and the regeneration extent at 72 hpa. As previously shown ^22^, Nox inhibition alone decreased both proliferation at 24 hpa and the regeneration extent at 72 hpa. Interestingly, Hh pathway stimulation with purmorphamine, a Smoothened agonist (Smo-A) that stimulates both early proliferation and increases the regeneration extent, completely rescued the effect of NOX inhibition on these two phenotypes (**Fig. 3A-A”**). Taken at face value, these results favor the interpretation that Hh signaling is downstream of NOX activity in blastema formation.

**Figure 3:**
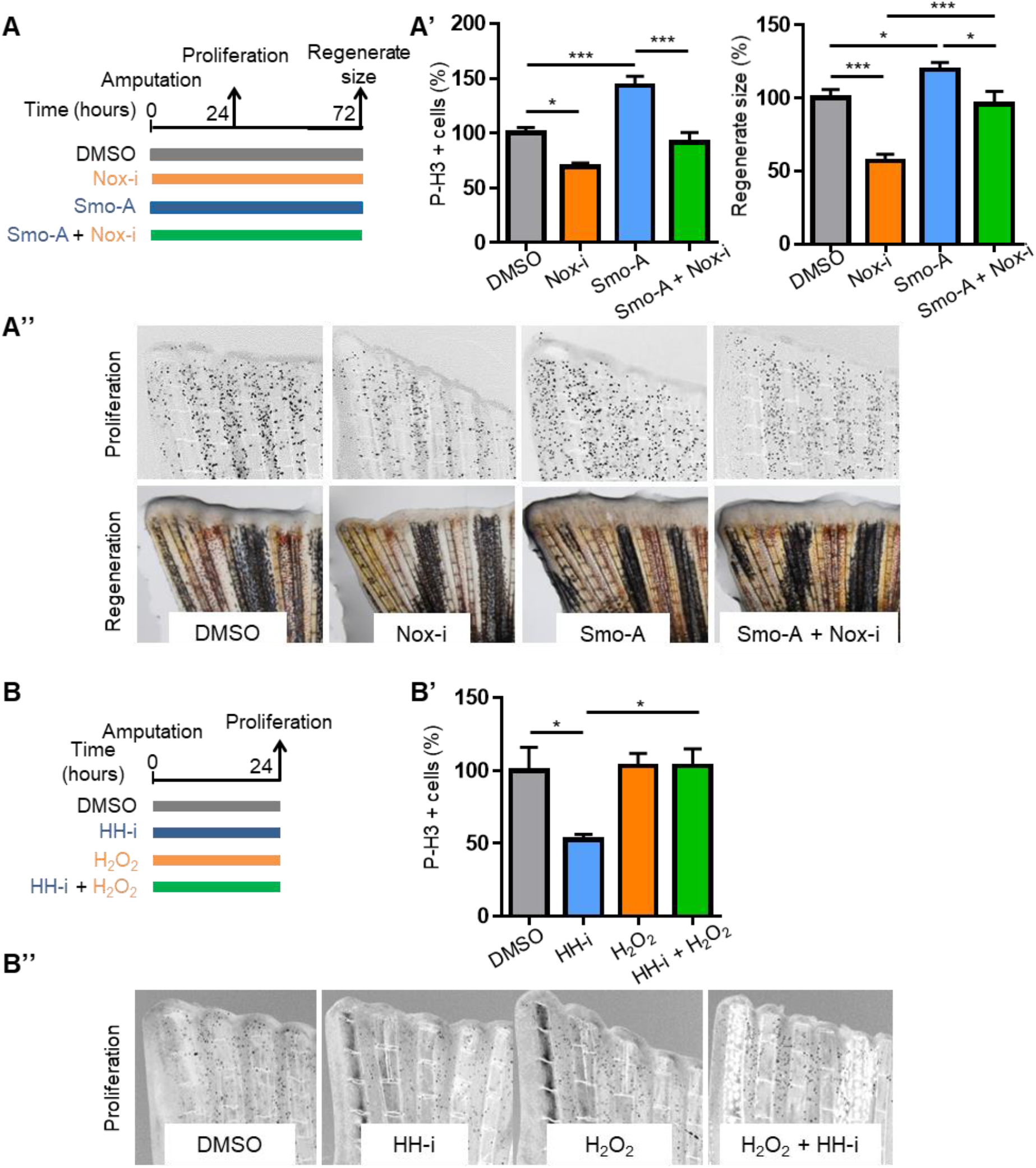
Crosstalk between the H_2_O_2_ level and the Hh pathway. (**A** and **B**) Schematics of the experimental procedures. (**A’** and **B’**) Proliferation in the epidermis at 24 hpa was monitored by measurement of the phosphorylated histone H3 (P-H3) level. Representative pictures are shown in A’’ and B’’. The efficiency of regeneration was quantified at 72 hpa. Representative pictures are shown in A’’. (**A-A’’**) Proliferation and regeneration were measured after NADPH oxidase inhibitor (Nox-i), purmorphamine (Smo-A) or vehicle (DMSO) treatment. (**B-B’’**) Proliferation was measured after cyclopamine (HH-i), hydrogen peroxide (H_2_O_2_) or vehicle (DMSO) treatment. The error bars indicate the SEM values (* p<0.05, ***p < 0.001).

We then performed a mirror experiment combining Hh pathway inhibition with H_2_O_2_ stimulation and measured the effect on stump cell proliferation. As previously shown ^22^, Hh inhibition alone was sufficient to significantly decrease proliferation at 24 hpa. Although H_2_O_2_ treatment alone had no effect by itself, its application in combination with Hh inhibition completely rescued the effect of Hh inhibition on proliferation (**Fig. 3 B-B”**). This pattern favors the interpretation that H_2_O_2_ signaling is downstream of Hh signaling in the same process, confirming that a regulatory loop indeed exists between these two signaling systems during the first 24 hours of adult fin regeneration.

### Early induction of Hh signaling components during regeneration

While the time course of ROS production over the first 24 hpa has previously been studied in some detail ^13^, no such analysis exists for Shh expression in this early phase, except for our previous observation that the *shha* locus is active very soon after amputation, as shown by GFP immunodetection in a shha:GFP transgenic line ^22^.

We thus performed real-time quantitative RT–PCR to characterize the time course of mRNA expression for major elements of the canonical Hh signaling pathway during the first 24 hpa, as a prerequisite to the study of the H_2_O_2_-Shh relationship (**Fig. 4**). In agreement with the early detection of GFP in our previous experiments ^22^, *shha* mRNA was already enriched above the background level at 3 hpa and increased until 24 hpa (with a plateau at 15–18 hpa). In addition, mRNAs of components of the pathway known to be targets of Hh signaling were studied. The mRNA levels of *ptch1* and *ptch2* increased significantly between 6 hpa and 15 hpa, as did that of *smo*, while the mRNA levels of the transcriptional effectors of Hh signaling slightly fluctuated below (*gli1*) or above (*gli2* and *gli3*) the baseline levels during the first 24 hpa. These results are consistent with the activity of some form of Hh signaling during the early module of fin regeneration. In addition, they point to a marked difference between the early and intermediate modules, with *ptch1* and *smo* specifically induced during the intermediate module, whereas *shha* activation starts earlier and appears more evenly distributed between the two modules.

**Figure 4:**
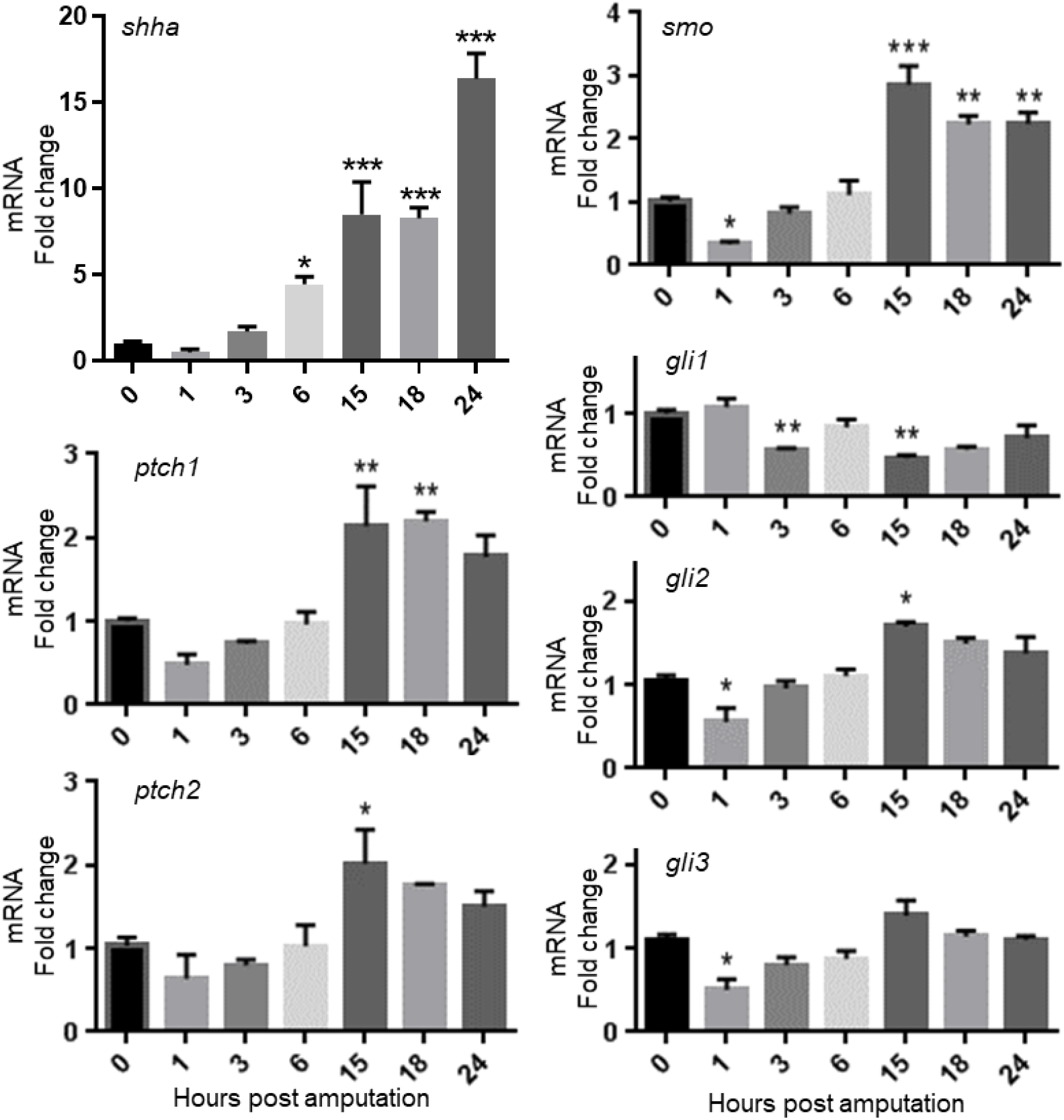
Early expression of Hh pathway members. Gene expression was analyzed through quantitative RT–PCR during adult fin regeneration. The level in the 0 hpa sample was set to 1. The error bars indicate the SEM values (* p<0.05, **p< 0.01, ***p < 0.001).

### The feedback loop between H_2_O_2_ and Hh signaling is already active during the early module

To test whether a reduction in the H_2_O_2_ level after amputation can modify the early activity at the *shha* locus (as we showed at 48 hpa, ^22^), we measured the abundance of GFP at 1 and 3 hpa in shha:GFP transgenic fish incubated with NOX-i (**Fig. S1 and Fig. 5A, B-B’**). This early inhibition was sufficient to reduce the level of GFP expression by the *shha* locus. We then examined the impact of NOX inhibition on the expression of *shha* and other Hh pathway components by real-time quantitative RT–PCR (**Fig. 5A, C**). Impairing NOX activity for 3 hours induced a significant reduction in *shha* expression but had no effect on *ptch1, ptch2, smo, gli1, gli2* or *gli3* expression. The reduction in shha expression was transient, since its protein and mRNA levels were not modified after 15 hours of treatment (**Fig. S2**).

**Figure 5:**
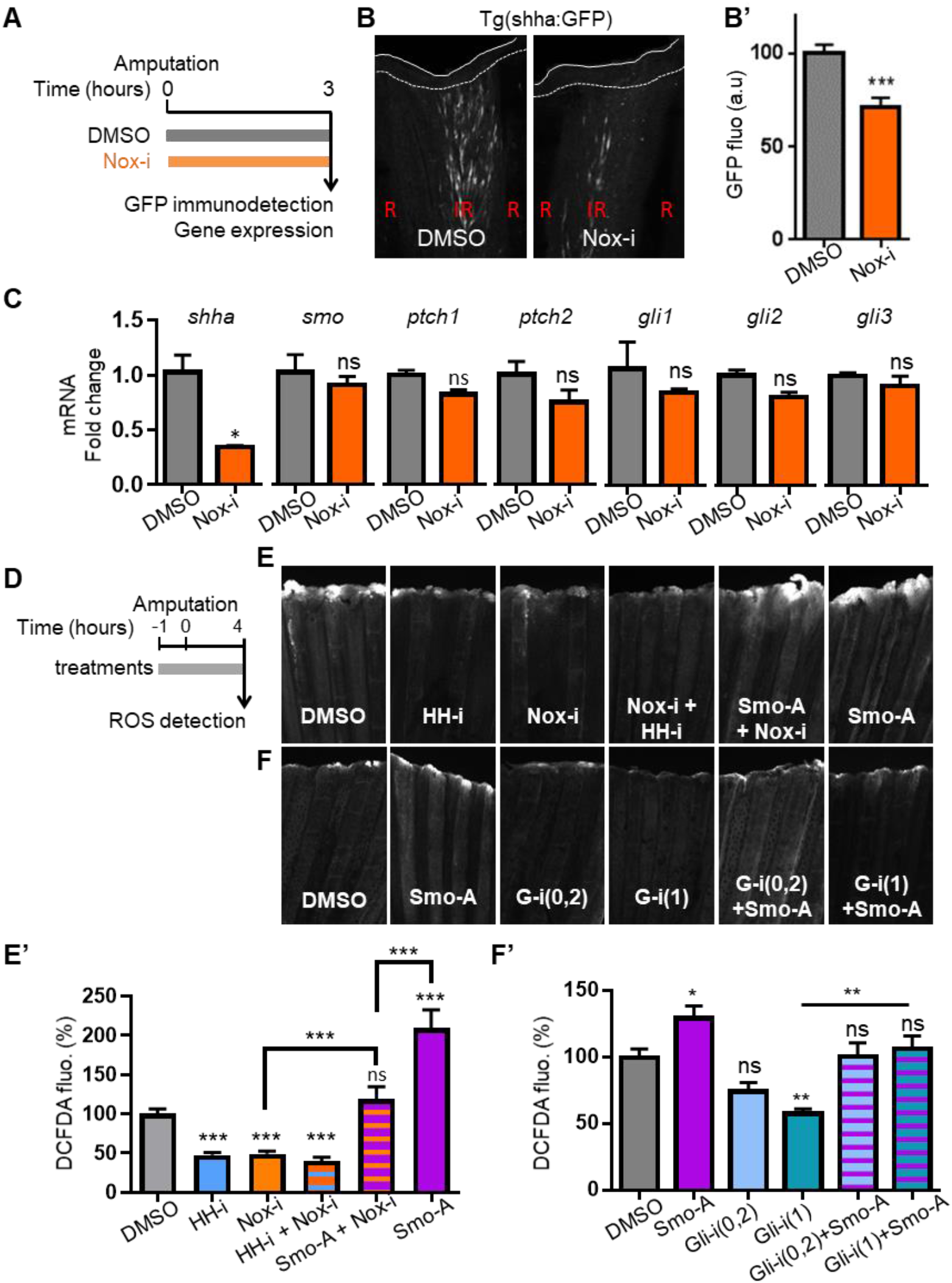
Early crosstalk between H_2_O_2_ and Hh signaling. (**A and D**) Schematics of the experimental procedures. (**B**) Cells expressing Shh were visualized at 3 hpa by immunodetection of GFP in shha:GFP transgenic fish after NADPH oxidase inhibitor (Nox-i) or vehicle (DMSO) treatment. Representative pictures are shown. Dotted line: amputation plane. Dashed line: distal part of the fin. R: ray; IR: inter-ray region. (**B’**) Quantification of GFP immunodetection. (**C**) Gene expression was analyzed by quantitative RT–PCR in the regenerated fin at 3 hpa after Nox-i or DMSO treatment. The level in the DMSO-treated sample was set to 1. (**E-E’**) ROS detection at the level of the amputation plane at 4 hpa after NADPH oxidase inhibitor (Nox-i), cyclopamine (HH-i), purmorphamine (Smo-A) or vehicle (DMSO) treatment. Representative pictures are shown in E’. (**F-F’**) ROS detection at the level of the amputation plane at 4 hpa after purmorphamine (Smo-A), Gli1/2 inhibitor (Gli-i at 0, 2 or 1 µM) or vehicle (DMSO) treatment. Representative pictures are shown in F’. The error bars indicate the SEM values (* p<0.05, ** p<0,001, ***p < 0.001).

To evaluate whether the interaction between H_2_O_2_ and Hh signaling is bidirectional also during this early phase, we modulated Hh signaling with HH-i or SmoA and analyzed ROS production at 4 hpa (**Fig. 5D, E-E’**). We verified that Hh signaling was necessary for ROS production beginning in the earliest phase: indeed, HH-i treatment strongly reduced the ROS level, while SmoA increased the ROS level. Fish were also cotreated with Nox-i and HH-i. Under this condition, no cumulative effect was observed, suggesting that Hh and redox signaling were in the same pathway. This hypothesis was confirmed by the rescue of Nox-i inhibition by SmoA treatment. Shh and H_2_O_2_ are thus part of a regulatory loop that is active even in the early module during fin regeneration. Finally, treatment of zebrafish subjected to fin amputation with the Gli-1/2 inhibitor (Gli-i) resulted in a ROS decrease similar to that obtained with HH-i, and this treatment was able to block the SmoA-dependent increase, providing evidence that Shh operates in this regulatory loop via a canonical pathway (**Fig. 5D, F-F’**).

## Discussion

A transient increase in the H_2_O_2_ level is required both for wound healing and slightly later to initiate the regeneration program in species ranging from flies to mice (for general review, see ^6^). However, the mechanisms controlling sustained H_2_O_2_ signaling specific to regeneration per se—as well as its timing, regulation and integration with other signaling pathways—are still imperfectly understood. This knowledge is a prerequisite for finding ways to improve regenerative processes, particularly in higher vertebrates.

The enzymes involved in H_2_O_2_ production have been examined in model species displaying very high regenerative capacities, i.e., fish and amphibians. It was first demonstrated that during the very early phase of wound healing in the zebrafish larval tail fin, a Duox-dependent gradient of H_2_O_2_ is generated within 3 min at the wound margin, peaks at 20 min and is required for the recruitment of distant leukocytes ^39^. Members of the NOX family were subsequently shown to be involved in regeneration per se, as is the case in the adult zebrafish caudal fin ^13,22^ and heart ^14^, gecko tail ^15^ and juvenile axolotl or Xenopus tadpole tail ^40,41^.

During the early response to injury, H_2_O_2_ is directly produced by Duox enzymes ^14,39^. Our results demonstrate that a transient decrease in Catalase activity, linked to a decrease in its gene expression, helps maintain a high level of H_2_O_2_. During the intermediate period, the sustained increase in the H_2_O_2_ level ^13,22^ is linked to elevated levels of SOD activity, which do not depend on changes in sod gene expression. This high level of H_2_O_2_ is subsequently reduced toward the end of the intermediate period by a marked increase in catalase activity, which does not depend on a change in catalase gene expression.

In parallel, Shh has been shown to play a role during the late phase of fin regeneration in zebrafish ^28–31,42,43^, tail regeneration in newts and Xenopus ^44,45^ and limb regeneration in axolotls ^46^. These studies looked at blastema growth and morphogenesis of the new appendage and were accordingly conducted later than 48 hpa. Recently, we discovered that Shha is expressed much earlier by activated Schwann cells, first migrating in the stump of the inter-ray region between 3 hpa and 12 hpa and then accumulating at the tip of the ray (between 24 hpa and 36 hpa) to reconstitute the Shh-positive cell population crowning the ray in uninjured fins ^22^. In this study, we characterized the early expression of several members of the Hh pathway during fin regeneration, and the dynamic profiles of expression observed during the first 24 hours suggested that Hh signaling is already active in the early module. This was shown in another series of experiments studying the feedback loop between Hh and H_2_O_2_ signaling (see below).

A link between Shh signaling and the redox status has previously been identified in different species and various pathological contexts. During mammalian heart injury, Hh signaling protects cardiomyocytes from H_2_O_2_-induced apoptosis ^47,48^. In rodent cortical neurons under oxidative stress, exogenous addition of Shh enhances SOD and glutathione peroxidase (GSH-PX) activity, protecting these cells from apoptosis ^34,49^. Hh signaling increases SOD activity in mouse astrocytes treated with molecules secreted by a parasite nematode ^50^. In Xenopus larvae, Shh regulates spinal cord and muscle regeneration through a noncanonical pathway ^51^. In zebrafish larvae, damage-induced ROS contribute to tail regeneration by repositioning Shh-producing cells ^52^. In addition, we discovered that physiological interactions between H_2_O_2_ and Hh signaling can be reciprocal, as is the case during both retinal projection development ^35^ and adult appendage regeneration ^22^.

Our present results clearly demonstrate the existence of a feedback loop between the two signaling systems during adult tail regeneration that is active beginning in the first 24 hours. On the one hand, Hh signaling affects the H_2_O_2_ level during the intermediate module by controlling SOD activity without changing *sod* gene expression by a mechanism that remains to be elucidated. On the other hand, the effect of inhibiting or increasing H_2_O_2_ signaling on regeneration is counteracted by activating or inhibiting, respectively, Hh signaling—and vice versa—within the first 3–4 hours after amputation. In addition, we found that Shh impacts H_2_O_2_ production via a Smo- and Gli-dependent canonical pathway. This is an interesting finding, since in a different biological system (HeLa cells), we recently identified a noncanonical pathway for Shh stimulation of H_2_O_2_ production, acting via the BOC-DOCK/ELMO-Rac1 axis ^36^.

The identity of the target in ROS signaling-mediated regulation of regeneration is still unknown. At 3 hpa, we observed that impairing NOX activity induced a significant reduction in shh expression but had no effect on the mRNA levels of other pathway components. The major action target of H_2_O_2_ is the reversible oxidation of sulfur in target proteins ^5^. Modulation of the H_2_O_2_ level might control the function of Hh pathway components or modify their expression by changing the oxidative status and thus the activity of relevant transcription factors. For example, Nkx2.1, which is involved in Shh expression in the ventral forebrain ^53^, belongs to the NK-2 homeoprotein family, whose members contain a conserved cysteine residue in the NK2-SD sequence. We described that Smo-dependent Hh signaling controls ROS levels during zebrafish fin regeneration. Smo is known to contain a highly conserved cysteine-rich domain crucial for its function and downstream Hh signaling ^54^. In a feedback loop, the oxidative status of this motif and the activity of this protein could be modulated by ROS levels. In addition, we observed that H_2_O_2_ impacts Shh trafficking and maturation during zebrafish development ^35,55^. This could also be the case in other contexts or species. Heparan sulfate proteoglycans (HSPGs), a class of proteins found on the cell surface and in the extracellular matrix, have been shown to regulate morphogen signaling, as is the case for Hh ^56^, particularly through their sulfation ^57^ (For review, see ^58^). For example, HSPGs facilitate Shh oligomerization and release ^59^. Among HSPGs, most glypicans and all syndecans are expressed in partially overlapping regions in the regenerating caudal fin ^60^. H_2_O_2_ was found to be able to modify heparan sulfate biosynthesis through transcriptional regulation of genes related to glycoconjugate metabolism in a keratinocyte cell line ^61^. These results suggested that ROS levels modify heparin sulfate chains and might have an effect on ligand distribution and signaling pathway activation.

Our demonstration of a feedback loop between Hh and H_2_O_2_ signaling also has broader implications, given the various crosstalk taking place during regeneration between Hh and other signaling pathways, such as that through BMPs, FGFs, or retinoic acid (RA) ^28,30,42,62,63^. This is particularly interesting in the case of Wnt signaling because several Wnt family members and pathway components are expressed during fin regeneration, and it was shown that the Wnt/β-catenin pathway regulates blastema proliferation through Hh signaling ^62^. However, most of this crosstalk between signaling pathways was studied after blastema formation, and it would be worth examining these interactions during the initial hours post amputation and how H_2_O_2_ might regulate this crosstalk.

More generally, cellular redox homeostasis and Hh signaling are involved in various cellular processes in physiological and pathological conditions, such as degenerative diseases, congenital syndromes and cancer (for review, see ^7,64,65^). Our results provide evidence for reciprocal interactions between Hh and H_2_O_2_ signaling that might be crucial in controlling these processes and provide insight into new treatments.

## Acknowledgments

The authors would like to thank Alain Prochiantz for constant support. This work received support under the program “Investissements d’Avenir” launched by the French Government and implemented by the ANR, with the following references: ANR-10-LABX-54 MEMO LIFE - ANR-11-IDEX-0001-02 PSL* Research University.

## Author contributions

C.R. and S.V. designed the experiment. M.T., R.M.D.S., M.A. and C.R. performed the experiments. C.R. wrote the manuscript. C.R., M.V. and S.V. analyzed the data and edited the manuscript.

