## Supplementary material for "An early Shh-H_2_O_2_ feedback loop controls the progression of the regenerative program during adult zebrafish fin regeneration": Table S1

Sup. Material for :

**Table S1: key resources table**

| Reagent type | Designation | Source | Identifier | Additional information (Final concentration) |
|---|---|---|---|---|
| Antibody | p-Histone H3 | Santa Cruz | Sc-8656R | Mitotic marker, (1:200) |
| | GFP | Abcam | ab13970 | For assessment of GFP-positive cells, (1:1000) |
| Kit | Catalase assay | Cayman Chemical | 707002 | CAT activity |
| | Superoxide dismutase assay | Cayman Chemical | 706002 | SOD activity |
| | Monarch total RNA miniprep | New England Biolabs | T2010S | Total RNA extraction, 50 preps |
| Chemical compound, drug | Cyclopamine | Calbiochem | 239803 | HH-i, 10 mM stock in DMSO (1 µM) |
| | GANT61 | Tocris | 3191 | Gli-i, 20 mM stock in DMSO (1 µM) |
| | Purmorphamine | Calbiochem | 540220 | Smo-A, 10 mM stock in water (1 µM) |
| | Hydrogen peroxide | Sigma–Aldrich | H1009 | $H_2O_2$, 10 mM stock in water (0.2 µM) |
| | Dimethyl sulfoxide | Sigma–Aldrich | D8418 | DMSO, solution |
| | VAS2870 | Enzo Life Sciences | BML-EI395 | Nox-i, 2.8 mM stock in DMSO (10 nM) |
| | 2’,7’-dichlorofluorescein diacetate | Calbiochem | 287810 | $H_2$DCFDA, 0.5 M stock in DMSO (33 nM) |
