## Supplementary material for "An early Shh-H_2_O_2_ feedback loop controls the progression of the regenerative program during adult zebrafish fin regeneration": Table S2

Sup. Material for :

**Table S2: Numbers of specimens used per experiment.**

| | | |
|---|---|---|
| **Fig. 1** | **B** | 0 h: 5, 1 h: 5, 15 h: 6, 30 h: 8, 72 h: 5. 4 fins per sample |
| | **C** | 0 h: 5, 1 h: 5, 15 h: 9, 30 h: 6, 72 h: 5. 4 fins per sample |
| | **D** | 0 h: 6, 1 h: 6, 3 h: 3, 6 h: 6, 15 h: 3, 18 h: 3, 24 h: 3. 4 fins per sample |
| | **E** | 0 h: 6, 1 h: 6, 3 h: 3, 6 h: 6, 15 h: 3, 18 h: 3, 24 h: 3. 4 fins per sample |
| **Fig. 2** | **A’** | DMSO—3 h: 3, 15 h: 4, 30 h: 4. HH-i—3 h: 3, 15 h: 4, 30 h: 4. 4 fins per sample |
| | **A’’** | DMSO—15 h: 4, 30 h: 4. HH-i—15 h: 4, 30 h: 4. 4 fins per sample |
| | **B’** | DMSO: 3, HH-i: 3. 4 fins per sample |
| **Fig. 3** | **A’** | Left DMSO: 13, Nox-i: 17, Smo-A: 12, Smo-A+Nox-i: 16 |
| | | Right DMSO: 14, Nox-i: 20, Smo-A: 29, Smo-A+Nox-i: 15 |
| | **B’** | DMSO: 6, HH-i: 7, $H_2O_2$: 7, HH-i+$H_2O_2$: 7 |
| **Fig. 4** | | 0 h: 6–9, 1 h: 3–6, 3 h: 3–6, 6 h: 4–9, 15 h: 3, 18 h: 3, 24 h: 3. 4 fins per sample |
| **Fig. 5** | **B’** | DMSO: 8, Nox-i: 13 |
| | **C** | 0 h: 3, 3 h DMSO: 3, 3 h Nox-i: 3. 4 fins per sample |
| | **E’** | DMSO: 76, HH-i: 83, NOX-i: 69, HH-i+NOX-i: 28, Smo-A+Nox-i: 31, Smo-A: 32 |
| | **F’** | DMSO: 41, Smo-A: 35, Gli-i(0,2): 42, Gli-i(1): 48, Gli-i(0,2)+Smo-A: 44, Gli-i(1)+Smo-A: 34 |

| | | |
|---|---|---|
| **Fig. S1** | **B** | DMSO: 5, Nox-i: 5 |
| **Fig. S2** | **B** | DMSO: 7, Nox-i: 8 |
| | **C** | DMSO: 3, Nox-i: 3. 4 fins per sample |
