## Supplementary material for "An early Shh-H_2_O_2_ feedback loop controls the progression of the regenerative program during adult zebrafish fin regeneration": Fig. S1

Sup. Material for :

**Figure S1: Early module**

**(A)** Schematic of the experimental procedure. **(B)** Quantification of GFP immunodetection at 1 hpa. Cells expressing Shh were visualized at 1 hpa by immunodetection of GFP in shha:GFP transgenic fish after NADPH oxidase inhibitor (Nox-i) or vehicle (DMSO) treatment. The error bars indicate the SEM values (***p < 0.001).

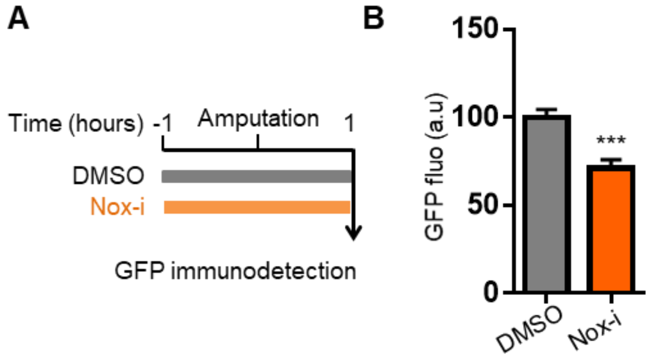
