## Supplementary material for "An early Shh-H_2_O_2_ feedback loop controls the progression of the regenerative program during adult zebrafish fin regeneration": Fig. S2

Sup. Material for :

**Figure S2: Intermediate module**

**(A)** Schematic of the experimental procedure. **(B)** Quantification of GFP immunodetection at 15 hpa. Cells expressing Shh were visualized at 1 hpa by immunodetection of GFP in shha:GFP transgenic fish after NADPH oxidase inhibitor (Nox-i) or vehicle (DMSO) treatment. **(C)** Gene expression was analyzed by quantitative RT–PCR in the regenerated fin at 15 hpa after Nox-i or DMSO treatment. The level in the DMSO-treated sample was set to 1. The error bars indicate the SEM values.

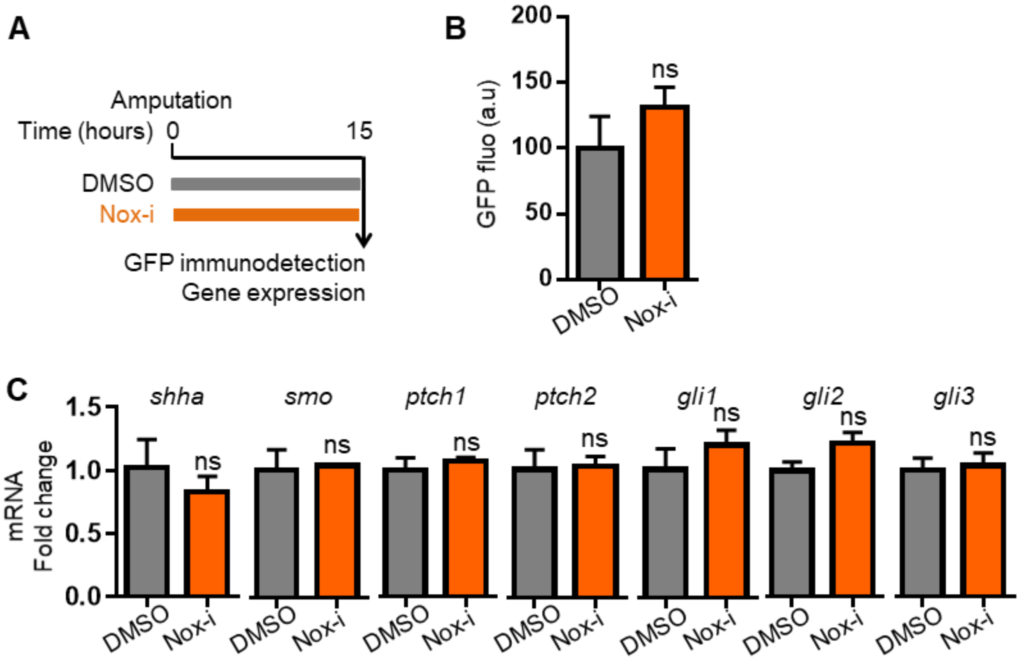
